# State-selective inhibition of alpha-synuclein aggregation by *de novo* oligomer-binding proteins

**DOI:** 10.64898/2026.07.13.738012

**Authors:** Yunxuan Zhang, Hannah L. Han, Lei Ortigosa-Pascual, Uri Z. Miles, Jiapeng Wei, Lexie Berkowicz, Finn Snow, Daizheng Tu, Georg Meisl, Shrutika Sansaria, Timonthy J. Nott, Heike Laman, Randal Halfmann, Andrew C. McShan, Danny D. Sahtoe, Tuomas P.J. Knowles

## Abstract

The intrinsically disordered, non-amyloid-β component domain of α-synuclein (αSyn) drives aggregation in Parkinson’s disease but lacks a stable epitope for structure-based drug design. We used deep learning-based protein design to generate *de novo* binders capturing this region in a β-strand conformation. Three designs bound αSyn with nanomolar affinities, with solution NMR spectroscopy supporting the intended fold. All three suppressed fibril formation at substoichiometric ratios, and fitting of aggregation kinetics resolved the microscopic steps inhibited by each. Notably, one binder preferentially engaged oligomers and inhibited secondary nucleation, both closely linked to toxicity. In cells, all three suppressed seeded αSyn assembly, with the oligomer binder being the most effective. Together, these findings establish a set of molecules that target distinct aggregation intermediates and mechanistically reshape αSyn assembly.

## Main Text

The misfolding and aggregation of α-synuclein (αSyn) into β-sheet rich amyloid fibrils is a defining molecular hallmark of Parkinson’s disease and related synucleinopathies(*1–3*). αSyn is intrinsically disordered in its monomeric form(*4*) but populates an extended β-strand conformation within mature fibrils, with the central non-amyloid β component (NAC, residues 61 to 95) forming the rigid amyloid core(*5–7*). Despite its central role in pathology, the NAC domain has remained a difficult therapeutic target because its lack of persistent structure deprives ligand discovery of a stable epitope used to identify selective small molecules or antibodies(*8–12*). Additionally, commercial reagents directed towards higher order species, such as oligomers or mature fibrils, do not intercept the initiating monomeric motif(*8*, *13*), which is the primary source of new nuclei and a principal driver of secondary nucleation(*14–16*).

Recent advances in deep learning-based protein design offer a route past this structural barrier. Diffusion-based generative models and sequence design networks can now produce compact scaffolds that recognise flexible and aggregation prone peptide motifs(*17–20*), and the concept has been demonstrated for amyloidogenic segments of amyloid β(*21*) and for disordered targets including amylin and G3BP1(*22*). However, target engagement alone does not establish control of aggregation, which requires resolving the species recognised and the microscopic steps that govern the aggregation of the full-length protein(*23*, *24*). These species - the soluble monomer, transient oligomer, or mature fibril - differ substantially in their toxicity and capacity to seed new aggregates(*25–27*).

In this work we design *de novo* protein binders against aggregation prone motifs within the αSyn NAC domain using the RFdiffusion/ProteinMPNN pipeline(*17*). The protocol yields compact, stable binders that engage the NAC domain with high affinity and, in the best case, strict selectivity against related amyloid cores, with solution NMR spectroscopy confirming the intended structural models. Combining microfluidic diffusional sizing (MDS) with thioflavin T (ThT) kinetics, we show that the binders arrest fibril formation substoichiometrically through distinct microscopic steps. They also suppress seeded assembly of αSyn in cells, with a potency ordering that follows from the microscopic assignment. We thereby move *de novo* binder design beyond target engagement toward the mechanism-guided inhibition of aggregation, establishing a generalisable route to amyloid cores long considered undruggable.

### Computational design of αSyn binders

We hypothesised that high affinity binders could be designed against αSyn fibril structures presenting the NAC domain in its β-strand configuration (Fig. 1a). Seven subregions within the NAC domain were selected as design targets(*7*) (fig. S1), and 30,000 backbones were generated using RFdiffusion(*17*). Structure-based filtering on defined secondary structure of proteins (DSSP)(*28*) and solvent-accessible surface area (SASA) scores removed backbones with poor packing and excessive loop and helix content. We then generated 24 sequences per backbone using ProteinMPNN(*18*), and filtered them with ColabFold(*29*) based on metrics such as pLDDT, pAE, and Cα-RMSD. Twenty-one final candidates were selected for experimental validation(*30*) (Fig. 1b).

**Fig. 1.**
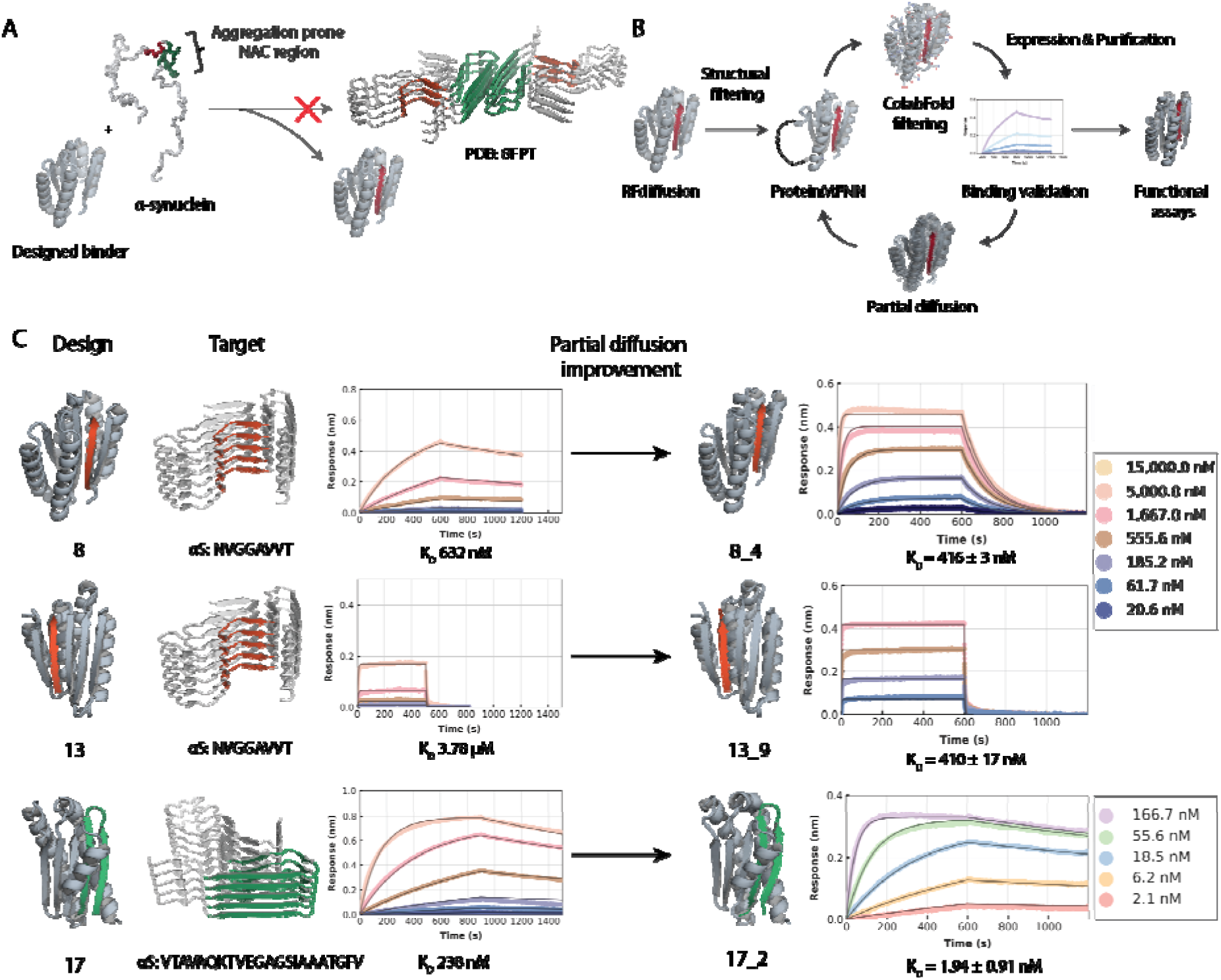
*De novo* design and affinity improvement of binders to the. α**Syn NAC core.** (**A**) Design concept. A *de novo* single chain scaffold is generated to complement and cap an exposed β-strand of the NAC region taken from the αSyn fibril, sequestering the segment into a stable, non-amyloidogenic complex. (**B**) The design pipeline starting with RFdiffusion2 for backbone design, followed by structural filtering on DSSP and SASA, sequence design with ProteinMPNN, and ColabFold filtering. (**C**) Left. Design models of three initial binders DFASYN008, DFASYN013 and DFASYN017 and their representative BLI sensorgrams. Right. Representative sensorgrams for the matured variants DFASYN008_4, DFASYN013_9 and DFASYN017_2. Error bars are derived from mean values ± standard deviation for three replicates (n = 3). Design models are coloured grey and target αSyn peptide red or green.

The 21 selected designs were expressed and screened for NAC-peptide binding by biolayer interferometry (BLI). Three of the designs (DFASYN008, DFASYN013 and DFASYN017) showed concentration-dependent binding, with dissociation constants of 632 nM, 3.78 µM, and 238 nM, respectively (Fig. 1c, left). The remaining 18 candidate designs gave no detectable response at any concentration tested.

To improve binding, partial diffusion was used to resample the backbone locally around each of the three successful binders, generating 500 new backbones per parent. Interface sequences were redesigned using ProteinMPNN under a bias favouring aromatic and aliphatic residues. Across designs that passed the same structural filters, the predicted metrics improved systematically relative to the first round (table S1). From each parent library, 5 to 10 matured variants were expressed and screened via BLI, and the tightest variant of each was carried forward as DFASYN008_4, DFASYN013_9 and DFASYN017_2, abbreviated 8_4, 13_9 and 17_2 (Fig. 1c, right, fig. S2). A single cycle of partial diffusion showed the most significant improvement for 17_2 of roughly 120-fold, from 238 nM to 1.94 nM.

### Designed αSyn binder adopts the expected structure in solution

To probe association of the designs with αSyn peptides in solution, U-[^15^N]-labelled 8_4, 13_9, and 17_2 were titrated with unlabelled αSyn peptides. Apo 8_4 and 13_9 gave 2D ^1^H-^15^N HSQC or HMQC spectra with excellent chemical shift dispersion, indicative of well-folded proteins (Fig. 2a, fig. S3). In contrast, the spectrum of apo 17_2 was of lower quality and lacked a number of the expected NMR peaks, suggesting conformational heterogeneity, dynamics, or insufficient folding. Addition of the αSyn peptide ^65^NVGGAVVT^72^ to 8_4 and 13_9 exhibited pronounced NMR peak sharpening, substantial chemical shift perturbations and the appearance of new peaks. Similarly, addition of the αSyn peptide ^74^VTAVAQKTVEGAGSIAAATGFV^95^ to 17_2 markedly improved spectral quality with the appearance of numerous new backbone amide resonances (Fig. 2a, fig. S3). These spectral changes suggest that peptide binding reduces conformational dynamics for all three binders and promotes a more ordered structural state, consistent with peptide occupation of the engineered binding pocket located between adjacent β-sheets.

**Fig. 2.**
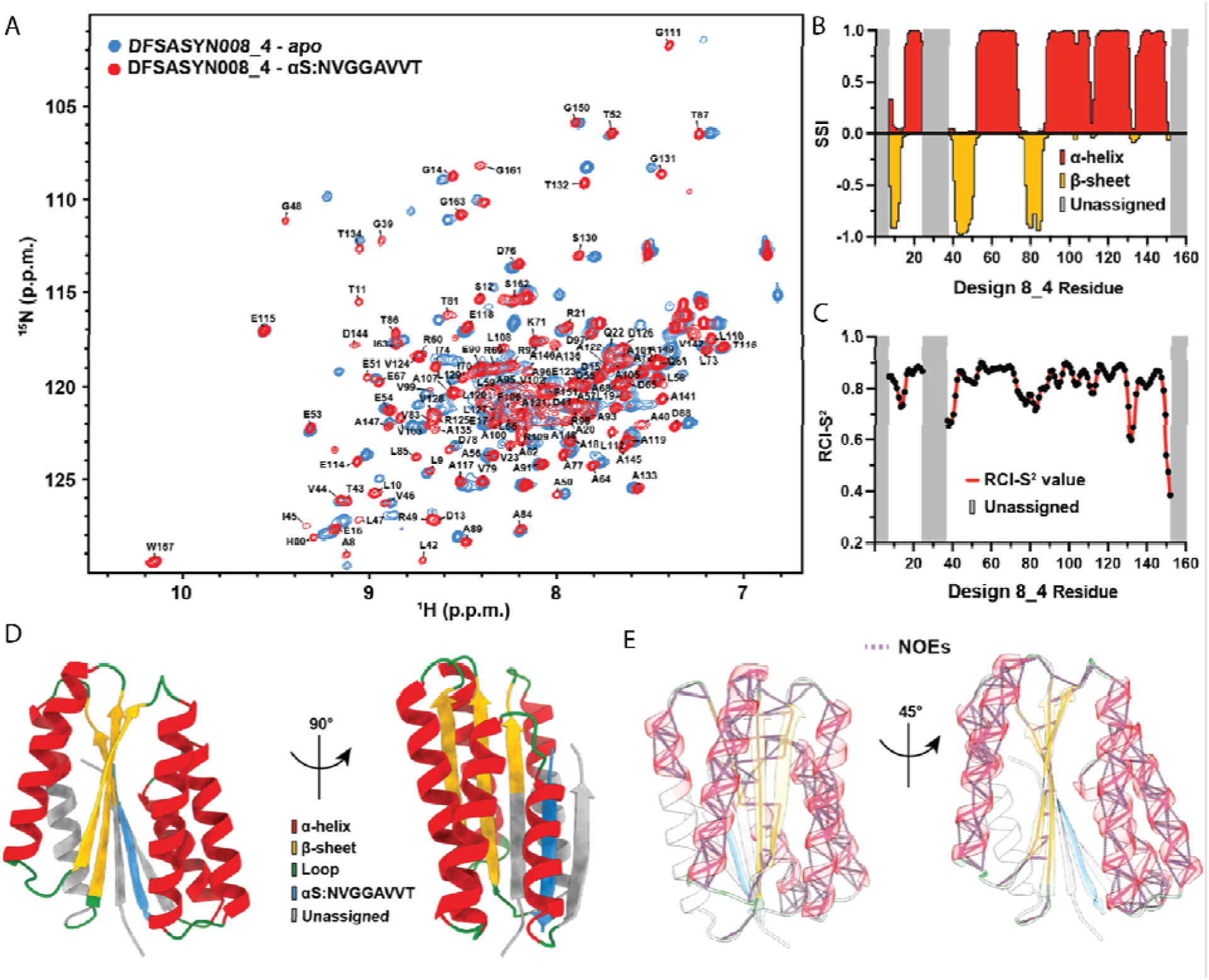
Solution NMR characterisation of the αSyn peptide/DFASYN008_4 complex. (**A**) 2D ^1^H-^15^N HSQC spectrum of U-[^15^N,^13^C] 8_4 in its apo state (blue) and αS: NVGGAVVT peptide bound state (unlabelled two-fold molar excess peptide, red) acquired at a ^1^H field of 800 MHz at 25 °C. (**B**) TALOS-N derived secondary structure index (SSI) as a function of 8_4 residue. β-sheets are marked in yellow, α-helices are marked in red, and unassigned zones are marked in grey. (**C**) TALOS-N derived random coil index order parameter (RCI-S^2^) as a function of DFASYN008_4 residue number. Residues with higher RCI-S^2^ values are more rigid; residue with lower RCI-S^2^ values are more dynamic. Unassigned regions are marked in gray. (**D**) Design model backbone for 8_4 coloured to match TALOS-N derived secondary structure in panel B. β-sheets are marked in yellow, α-helices are marked in red, loops are marked in green, αS: NVGGAVVT peptide marked in blue, and unassigned residues are marked in grey. (**E**) Through-space amide to amide NOEs (dotted purple lines) derived from 3D SOFAST HMQC NOESY experiments plotted onto the design model backbone for 8_4. Unassigned residues are coloured white.

The spectral quality of the NVGGAVVT/8_4 complex enabled assignment of 76.7% of the backbone resonances (H_N_, N_H_, Cα, Cβ, and CO) using standard sequential assignment strategies based on triple-resonance NMR experiments (Fig. 2a). Unfortunately, resonances corresponding to residues on the β-strands proximal to the peptide-binding groove were not observable, presumably due to unfavourable conformational exchange on the NMR timescale due to high off rate (Fig. 1c). Because of gaps in the assignments, the full three-dimensional NMR structure could not be determined. However, TALOS-N calculated secondary structure index (SSI) and random coil index order parameter (RCI-S^2^) values derived from assigned NMR chemical shifts provide direct evidence supporting the expected structure of 8_4’s design model (Fig. 2b-d). TALOS-N derived secondary structure elements for α-helices, β-sheets, and loops match well to the design model (Fig. 2b, c). Low RCI-S^2^ values for the assigned β-strands adjacent to the predicted peptide-binding site indicated increased backbone flexibility in these regions, consistent with their proximity to the peptide-binding interface (Fig. 2c). Through-space nuclear Overhauser effect (NOE) contacts between backbone amide protons in 8_4 were consistent with the predicted secondary structure, providing additional evidence that the NVGGAVVT/8_4 complex adopts the intended fold. Inter-β-strand NOEs further supported the presence of the designed antiparallel β-sheet (Fig. 2e). Overall, these results demonstrate that 8_4 adopts the intended secondary and tertiary structure upon binding the αSyn peptide NVGGAVVT.

### Binder selectivity across related amyloidogenic proteins

We profiled the three matured binders by BLI against peptides representing the aggregation cores of tau(*31*), amylin(*32*, *33*) and amyloid β(*34*) (table S2) to test for cross-specificity. 17_2 was strictly selective, showing no detectable binding to any of the three off-target peptides at the concentrations tested while K_D_ to its αSyn target was 1.94 nM (fig. S4). We attribute this selectivity to the design’s mode of engagement, since 17_2 caps a β-hairpin through a multi-point interface that includes a flexible C-terminal tail, a geometry that demands shape complementarity that the other amyloid cores do not provide. The two remaining binders were less discriminating. 8_4 retained a three to twentyfold preference for αSyn, whereas 13_9 bound all three target peptides more tightly than αSyn, most likely due to an open-faced architecture that presents a largely hydrophobic surface accommodating amyloidogenic cores non-specifically.

### Engagement across the aggregation states

We next asked how the binders engage full-length αSyn across its aggregation states using MDS, which reports binding as a change in hydrodynamic radius R_h_ when a labelled binder associates with other species(*35*). Monomer and oligomer affinities were measured on a Fluidity One-M platform suited to the 1 to 20 nm range, and fibril affinities on a custom single molecule platform (smMDS) that resolves large, slowly diffusing species and to deconvolute the bound and free fractions(*36*) (Fig. 3a,b, fig. S5). Kinetically trapped oligomers prepared by a previously established method(*37*) served as the proxy to the oligomeric species. Because binder affinity for fibrils can be polymorph-dependent, fibrils were formed under the same conditions used for the aggregation kinetics experiments.

**Fig. 3.**
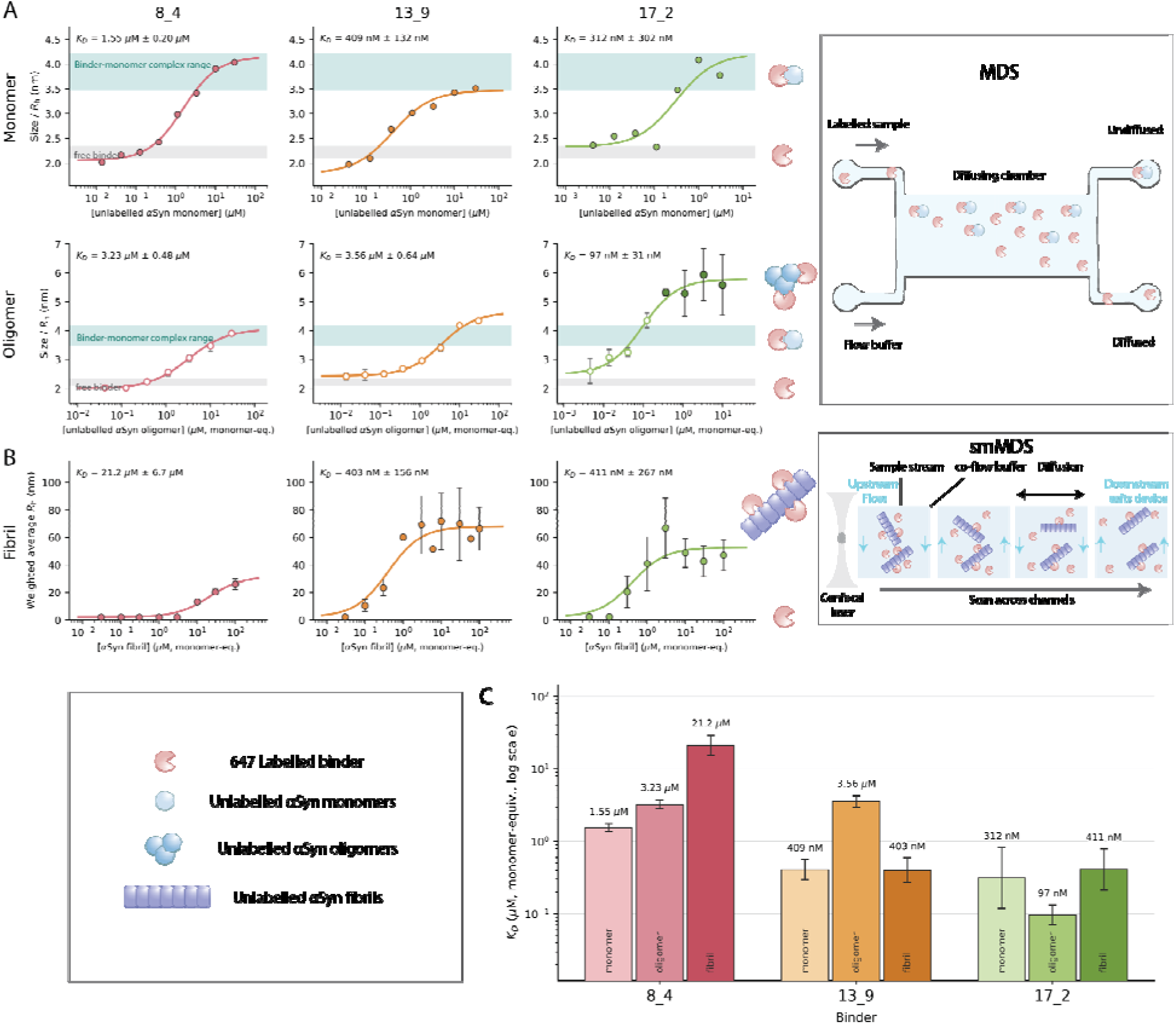
Binder selectivity for different. α**Syn aggregation species.** (**A**) (Left) Binding isotherm for 8_4, 13_9 and 17_2 against monomeric and oligomeric αSyn monitored through measurements of the size increase as the bound complex forms. (Right) Principles of the Fluidity One-M MDS measurement device. (**B**) Left: Binding isotherms for the binders against fibrillar αSyn. Right: Principles of smMDS to capture diffusion of larger sized species. (**C**) Summary of the fitted dissociation constants by species on a logarithmic scale. Error bars are derived from mean values ± standard deviation for three replicates (n = 3).

The binders engaged the full-length monomer more weakly than the isolated NAC peptide, as expected from the added steric context of the intact protein(*38*), with dissociation constants of 1.55 µM for 8_4, 409 nM for 13_9, and 312 nM for 17_2 (Fig. 3c). Against oligomers, complex size is also informative, as the monomer titrations define the complex size of a binder-monomer complex, allowing larger complexes to be assigned to oligomers. 8_4 never exceeds that size, so its apparent oligomer binding cannot be distinguished from monomer binding. 13_9 formed slightly larger complexes but bound ninefold weaker than the monomer, at 3.56 µM. Only 17_2 formed complexes clearly larger than a binder-monomer pair, and did so with the tightest affinity measured at 97 nM, threefold tighter than for the monomer. It is therefore the only binder that unambiguously engages oligomers, notable because small, soluble oligomers are strongly linked to toxicity(*25*, *26*). A preference of this kind can be explained for 17_2, which was designed against a β-hairpin, an arrangement of two strands that the monomer samples only transiently but that may be partly formed in the oligomer, where clustering of αSyn molecules increases the local concentration of compatible binding sites and thereby enhances apparent affinity.

Against fibrils, 13_9 and 17_2 bound comparably, at 403 and 411 nM, while 8_4 was far weaker at 21.2 µM. These are bulk averages over the whole fibril and thus report mainly on the lateral surface rather than on the fibril ends. Imaging gold-conjugated binders bound to fibrils by transmission electron microscopy (TEM) confirmed that both 8_4 and 17_2 engage fibrils directly, with 8_4 most often seen at the fibril ends and 17_2 along the sides (fig. S6). 8_4 therefore gave the clearest fibril end localisation despite the weakest bulk affinity. Thus, we reasoned that a whole-fibril dissociation constant reveals little about where a binder acts. We used aggregation kinetics below to resolve their functional engagement at the fibril ends and the lateral surfaces separately.

### Binders inhibit aggregation substoichiometrically through distinct microscopic steps

We next tested whether the binders inhibit αSyn fibril formation *in vitro* using ThT fluorescence assay. At 10% of binder concentration relative to 80 µM αSyn, all three binder showed complete inhibition of fibril formation for the full 180 h of the assay (Fig. 4b). Sequestration of bulk monomer cannot account for this, since it would require at least equimolar concentrations binder. Removing the equivalent 10% of monomer from an αSyn-only reaction (80 µM to 72 µM) produced almost no measurable delay, confirming that the binders must be acting on the aggregated or nucleating species of αSyn.

**Fig. 4.**
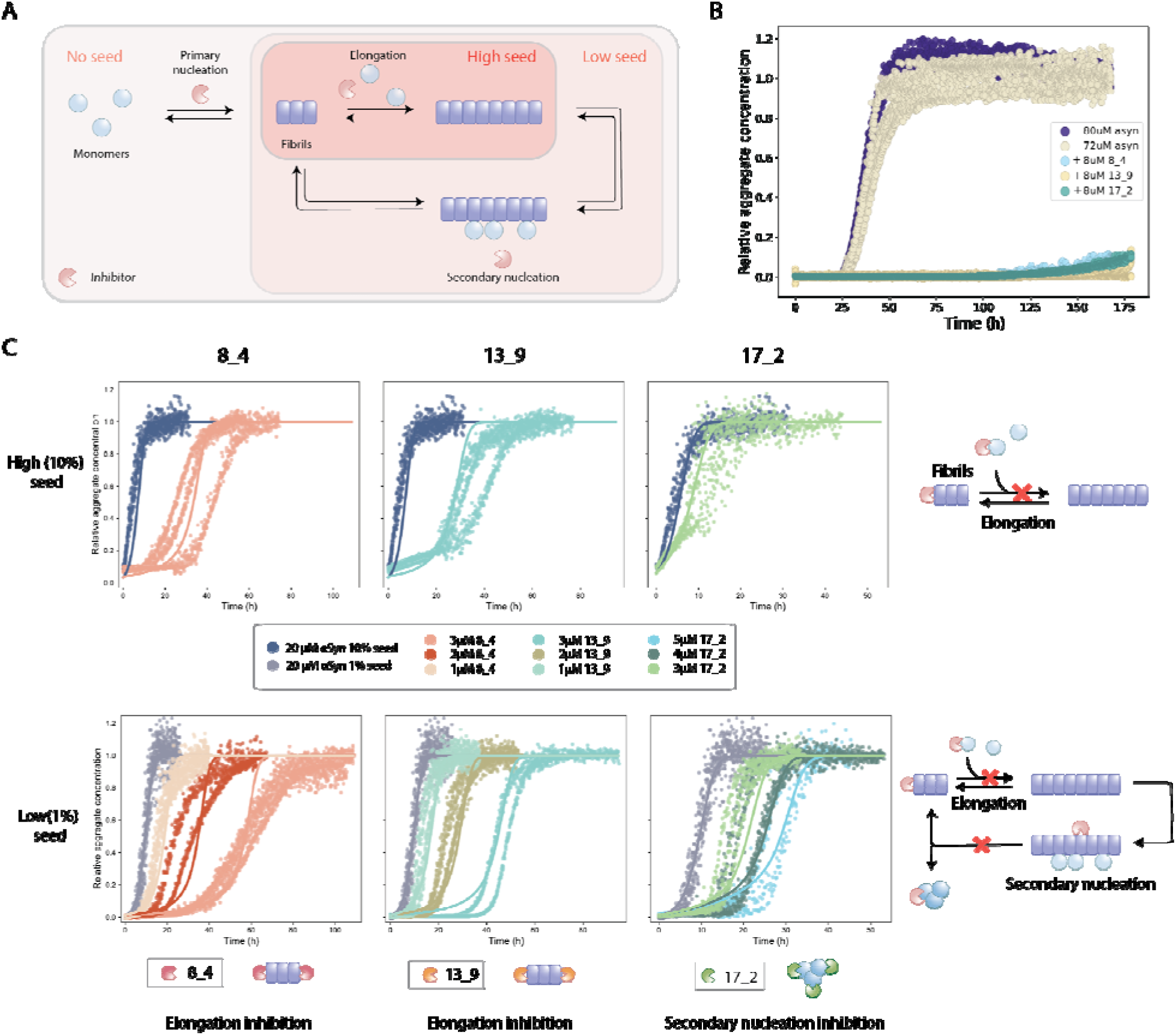
Effect of binders on. α**Syn aggregation kinetics.** (**A**) Microscopic steps of amyloid aggregation and the effect of seeds. At no seed all steps operate, at low seed primary nucleation is bypassed to report secondary nucleation and elongation, and at high seed elongation dominates. (**B**) Test for monomer sequestration. An 80 µM αSyn reaction was run in parallel with 10% (8 µM) of each binder, and a control experiment of αSyn only, removing 10% of the monomer to give 72 µM. (**C**) Binder titration with high seed (10%) and low seed (1%) conditions. Points are ThT data averaged across n=3 repeats. Global kinetics fitting were conducted on high and low seed conditions to investigate the binder contribution in elongation and secondary nucleation steps.

To identify which microscopic steps are affected, we ran seeded reactions of 20 µM αSyn at low (1%) and high (10%) seed concentrations, separating the contributions of secondary nucleation and elongation(*15*, *24*, *39*). Seeding bypasses primary nucleation. At low seed concentrations, the reaction remains autocatalytic and both secondary nucleation and elongation contribute, whereas at high seed concentrations the abundance of fibril ends renders secondary nucleation negligible and elongation dominates (Fig. 4a, fig. S7). Binder-free reactions were first fitted globally to a secondary-nucleation-dominated model using AmyloFit(*24*), and the resulting rate constants are held fixed (table S4). Binder containing reactions were then fitted globally to a model in which a binder may occupy either the fibril ends, with dissociation constant K_IE_, or the catalytic surface at which secondary nucleation proceeds, with dissociation constant K_IS_ (Fig. 4c, fig. S8). Because the binders are added at substoichiometric ratios and are depleted by the aggregates they bind, we did not assume that free binder remains in excess for kinetic analysis (fig. S9).

For 8_4 and 13_9, K_IE_ lies three and two orders of magnitude below K_IS_ (table S4), suggesting both binders as elongation inhibitors. This accounts for the strong inhibition seen at high seed and, because secondary nucleation is also fed by fibril mass, for the extended lag phase at low seed. For 8_4, this is also supported by the prominent end localisation seen in TEM (fig. S6). 13_9 binds the fibril tightly in bulk yet inhibits secondary nucleation only weakly, indicating that it can occupy the lateral surface of the fibril without specifically blocking the sites at which nucleation occurs. Since both binders were designed against a single short β-strand within the NAC core, a monomer docked at a growing end is templated into exactly this conformation, making the end a high-affinity site which a bulk measurement cannot resolve. Consistent with action at a scarce, rate-limiting site, titrating 8_4 under even higher (20%) seed gave near-complete suppression at a 20% binder ratio (fig. S10).

17_2 inhibits aggregation by an alternative route. Its K_IS_ of 127 nM is roughly 250-fold below its K_IE_, indicating potent inhibition of secondary nucleation with only a small effect on elongation. Its K_IS_ agrees within error with the 97 nM binding affinity against oligomers measured by MDS. We therefore assign 17_2 as capturing nascent oligomers as they are generated at the fibril surface and diverting them into a non productive complex, so that they do not mature into elongation competent fibrils. This sets 17_2 apart from the other two binders by inhibition mechanism, as it suppresses the autocatalytic cycle by engaging the species most closely linked to toxicity(*25*, *26*, *37*).

### Binders engage αSyn and suppress its seeded assembly in cells

Having resolved which steps of the aggregation pathway the binders inhibit *in vitro*, we asked whether target engagement and inhibition are retained in cells. We first constructed a recruitment assay in HeLa cells in which the αSyn NAC domain was anchored to the inner face of the plasma membrane as a fusion to mScarlet and the pleckstrin homology domain of phospholipase Cδ (39), and each binder was expressed as an EGFP fusion under inducible control (Fig. 5a). All three binders colocalised with the target, with Pearson coefficients of 0.57 for 8_4 and 0.77 and 0.90 for 13_9 and 17_2, respectively (Fig. 5b). Without target the same binders remained diffusely cytoplasmic (r = 0.05), and no colocalised signal appeared in the absence of doxycycline or of target transfection (fig. S11). The binders therefore locate the NAC motif specifically within a cellular context.

**Fig. 5.**
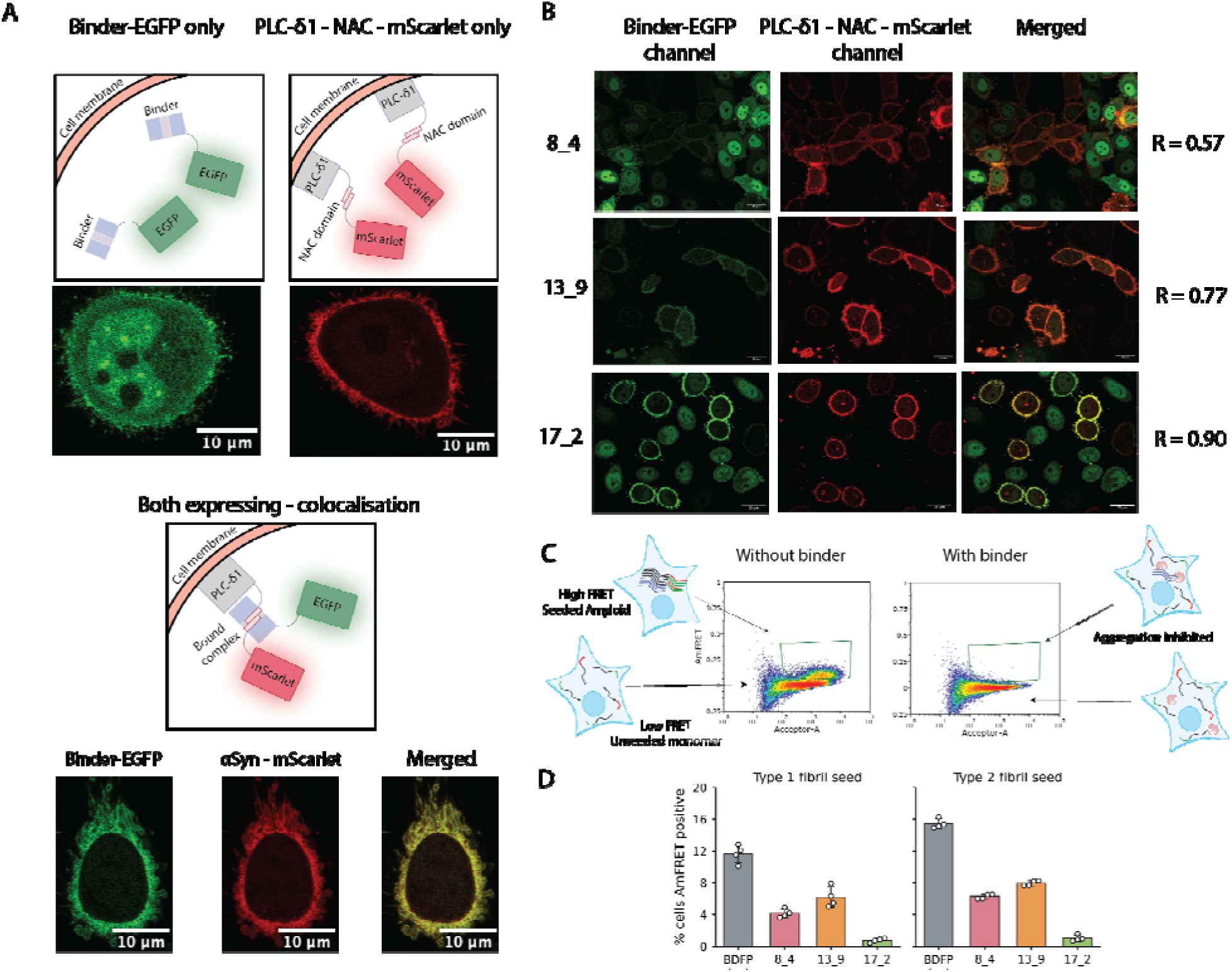
Engagement and inhibition of. α**Syn assembly in cells.** (**A**) Design of the recruitment assay. The NAC target is expressed as a fusion to mScarlet and the pleckstrin homology domain of phospholipase Cδ, anchoring it to the inner plasma membrane, while each binder is expressed as an EGFP fusion under doxycycline inducible control. (**B**) Live cell confocal imaging of the three binders. Columns show the EGFP binder channel, the αSyn mScarlet channel and the merged channels. Scale bars 20 µm. (**C**) Schematic of the DAmFRET experiment. Each data point represents a single cell, and those in which αSyn has aggregated give a high AmFRET signal, shown without binder (left) and with binder (right). (**D**) Percentage of cells within th AmFRET positive gate for each binder and both seeding polymorphs. Bars show the mean of n=4 repeats.

To further test whether the binders prevent full length αSyn assembly in cells, we used DAmFRET, a fluorescence resonance energy transfer (FRET)-based assay which scores αSyn aggregation in individual cells(*40*, *41*). HEK293T cells expressing mEos3.2 tagged αSyn A53T were seeded with 0.25 µg of preformed αSyn fibrils of two structurally distinct polymorphs, Type 1 or Type 2 (fig. S12), with each binder co-expressed. All three binders reduced the AmFRET-positive population under both polymorphs. 8_4 and 13_9 gave partial inhibition, between 47 and 64 %, whereas 17_2 lowered the AmFRET positive fraction by 94 % to a level indistinguishable from its unseeded baseline, indicating that seeding was abolished rather than delayed (Fig. 5c). Inhibition was closely matched for the two polymorphs, indicating action on a feature of the NAC core that they share, and was not accounted for by differences in target or binder expression.

The order of potency differed from high-seeded reactions *in vitro*, where an excess of fibril ends favours elongation inhibitors. Because intracellular seed availability is limited and aggregate amplification is required to cross the AmFRET threshold, the cellular assay instead favours inhibition of secondary nucleation, consistent with the strong activity of 17_2.

## Discussion

Deep learning-based protein design has only recently made intrinsically disordered and amyloidogenic sequences accessible to ligand discovery(*20–22*). Building on this, we designed *de novo* proteins that engage the NAC aggregation core of αSyn, obtaining binders with nanomolar affinity and, in the best case, strict selectivity over other amyloid cores. Binders raised against different NAC epitopes diverged in both the aggregation states they bound preferentially and the microscopic step they inhibited. 8_4 and 13_9 recognised a single short β-strand and they acted through capping the fibril ends and blocking elongation. Whereas 17_2, designed against a longer β-hairpin, instead binds oligomers and suppresses secondary nucleation. The choice of epitope within a fibril core may therefore determine which microscopic step a binder inhibits, raising the possibility that mechanism can be specified at the design stage rather than discovered afterwards. Since the only structural input required is a fibril, the approach should extend to other amyloids and, as polymorph structures accumulate, to polymorph-specific targeting.

Functionally, all three binders suppress fibril formation at substoichiometric ratios through a mechanism more specific than depletion of the bulk monomer pool. For 17_2, the dissociation constant fitted from the aggregation curves agrees within error with the affinity measured against isolated oligomers. It is also the most effective binder in cells, where amplification of a small amount of seed is dependent on secondary nucleation. Together, these measurements identify 17_2 as an oligomer-preferring inhibitor of secondary nucleation. Because secondary nucleation is the principal source of toxic αSyn oligomers(*16*, *26*, *42*), 17_2 should limit their generation, although a direct effect on oligomer levels and toxicity remains to be tested. Beyond their use as inhibitors, selective and state-resolved binders may also provide tools for interrogating and capturing defined aggregation states.

Parkinson’s disease still has no disease-modifying therapy, and the most advanced αSyn antibodies have yet to meet their efficacy endpoints(*43–45*), in part because immunisation raises antibodies against the charged termini rather than the hydrophobic NAC core that drives aggregation(*46*). A modality that engages the toxic oligomeric species directly is therefore attractive, and designed proteins carry practical advantages, being smaller, more stable, and genetically encodable(*47*). Although our data demonstrate specific target engagement and potent inhibition of aggregation *in vitro*, extending this to neuronal and *in vivo* models of spreading pathology is the important next step. Likewise, evaluating delivery across the blood–brain barrier and addressing the broader complexity of disease pathology will be key considerations for therapeutic translation. Even so, we have shown that binders to an undruggable aggregation core can be made selective for rare, rate-limiting intermediates, and can suppress aggregate proliferation both potently and specifically. This establishes *de novo* designed proteins as a credible strategy against amyloid formation, and as new tools for dissecting it.

## Acknowledgments

We would like to thank B.I.M. Wicky and L. Milles at the Institute for Protein Design for providing Golden Gate cloning vectors, N. Lorenzen for general discussion on drug development, and E. Axell for general discussion on amyloidosis and neurodegenerative diseases. Claude AI was used to assist with manuscript editing and with code for figure plotting.

## Funding

European Research Council grant DiProPhys 101001615 (TPJK) Stichting Parkinsonfonds (DDS)

Stowers Institute for Medical Research (RH) Novo Nordisk studentship (HLH)

Wilkinson Foundation studentship (YZ)

## Author contributions

Conceptualization: YZ, HLH, DDS Methodology: DDS, HLH

Investigation: YZ, HLH, LOP, LB, FS, DT, SS, UZM, ACM

Formal analysis: JW, GM Visualization: YZ, HLH

Funding acquisition: TPJK, DDS, RH Supervision: TPJK, DDS

Writing – original draft: YZ, HLH

Writing – review & editing: YZ, HLH, LOP, DDS, LB, JW, GM, DS, DT, TJN, HL, RH, ACM, TPJK

## Data, code, and materials availability

All data are available in the main text or the supplementary materials

## Supplementary Materials

Materials and Methods

Supplementary Text

Figs. S1 to S12 Tables S1 to S4

References (*48–62*)

