## Supplementary materials for "State-selective inhibition of alpha-synuclein aggregation by *de novo* oligomer-binding proteins"

Science
AAAS

Supplementary Materials for

### State-selective inhibition of alpha-synuclein aggregation by *de novo* oligomer-binding proteins

Yunxuan Zhang[1,2]†, Hannah L. Han[1,2]†, Lei Ortigosa-Pascual[2,8], Uri Z. Miles[5], Jiapeng Wei[1,2], Lexie Berkowicz[6,10], Finn Snow[7], Daizheng Tu[1,2], Georg Meisl[2,9], Shrutika Sansaria[10], Timonthy J. Nott[8], Heike Laman[7], Randal Halfmann[6,10], Andrew C. McShan[5], Danny D. Sahtoe[3,4]*, Tuomas P.J. Knowles[1,2,11]*

**The PDF file includes:**

Materials and Methods
Figs. S1 to S12
Tables S1 to S4
References (*48–62*)

#### Materials and Methods

*De novo* binder design

Binder backbones were generated using RFdiffusion2(*17*) targeting the Non-Amyloid Component (NAC) region of αSyn (PDB: 8FPT). A total of 30,000 backbones were generated against seven different target strands (Supplementary Fig. 1). To enrich scaffolds with favorable foldability and solubility profiles prior to sequence design, backbones were filtered based on secondary structure content (assigned using DSSP(*28*)) and solvent-accessible surface area (SASA).

Amino acid sequences were designed onto the filtered backbones using ProteinMPNN(*18*). For each backbone, 24 sequences were sampled, with cysteine and methionine residues explicitly excluded from the design process. Structural validation was performed using ColabFold (v1.5.2)(*29*, *48*), generating five models per design. Designs were filtered to meet all of the following thresholds: (1) high local structural confidence (pLDDT > 92); (2) high interface confidence (mean pAE < 5 and max pAE < 25); and (3) high structural fidelity to the design model (Cα-RMSD < 1.2 Å). Following visual inspection of interface packing and electrostatics, 21 designs were selected for chemical synthesis and experimental characterisation.

Binder optimisation via partial diffusion

Lead binder scaffolds were refined using the partial diffusion implementation of RFdiffusion (v1.1.0)(*17*). The diffusion process was initiated at a partial noising step of T=20 to allow for whole backbone fluctuations. Following backbone refinement, sequence optimisation was conducted using ProteinMPNN(*18*). The design process utilized a position-specific bias favoring aromatic and hydrophobic residues at the binding interface while penalizing polar and charged residues to optimize the desolvation profile of the interface.

Protein expression and purification

Synthetic genes obtained from Integrated DNA Technologies were cloned into custom expression vectors(*49*) via Golden Gate assembly with a C-terminal SNAC-GST-hexahistidine ($His_6$) tag(*50*). Peptide sequences for BLI were expressed with ubiquitin-AviTag-$His_6$ and Pro-Ala-Ser linker on the N-terminus.

Proteins were expressed and purified as previously reported(*21*, *30*). Briefly, vectors were transformed into BL21 (DE3) *E. coli* cells and expressed using TBII autoinduction media. Proteins were grown for 6 h at 37°C before 20°C overnight under antibiotics selection. Cells were isolated by centrifugation at 4,000×*g* and resuspended in lysis buffer (20 mM Tris, 100 mM NaCl, 50 mM imidazole, pH 8.0) containing protease inhibitors and bovine pancreas DNAse I. The cells were lysed by sonication, and the clarified lysate was incubated with nickel NTA beads for 1 h before washing with 10 column volumes of lysis buffer, 10 column volumes of high salt buffer (20 mM Tris, 1 M NaCl, 50 mM imidazole, pH 8.0), 10 column volumes of lysis buffer. The uncleaved proteins can be eluted with 7.5 mL of elution buffer (20 mM Tris, 100 mM NaCl, 500 mM imidazole, pH 8.0). For SNAC cleavage, the proteins were further washed with 10 column volumes of SEC buffer (1× PBS, pH 7.4) and 10 column volumes of SNAC cleavage buffer (100 mM

CHES, 100 mM Acetone oxime, 100 mM NaCl, 500 mM GnCl, pH 8.6). The beads were incubated overnight in 10 column volumes of SNAC cleavage buffer supplemented with 2 mM $NiCl_2$.

For mutated designs with free cysteines, 1 mM TCEP was added to every step of the purification process. After SNAC cleavage, the cleaved proteins were buffer exchanged into 1× PBS and labelled with fivefold excess of Alexa647-C2-maleimide and 1 mM TCEP for 3 h at room temperature.

The eluted and labelled proteins were all subjected to a SEC purification on Superdex 75 Increase 10/300GL columns (Cytiva) using 1× PBS as a final step of the purification process.

Peptide synthesis

Peptides for NMR were ordered and synthesized via Genscript at 99% purity.

Circular dichroism spectroscopy

Circular dichroism spectra were recorded on a Chirascan spectrometer (Applied Photophysics) in a cuvette with a 1 mm path length at a protein concentration between 0.4 and 0.6 mg/mL. Experiments were performed in 20 mM sodium phosphate pH 8. Spectra were acquired from 190 to 250 nm with a 1 nm step size.

Solution NMR spectroscopy

Uniformly $^{15}$N-labelled (U-[$^{15}$N]) 8_4, 13_9, and 17_2 were expressed in 1× M9 minimal media containing 1 $gL^{-1}$ $^{15}NH_4Cl$, 3 $gL^{-1}$ glucose, 0.1 $gL^{-1}$, $^{15}$N ISOGRO, and 12.5 $mgL^{-1}$ Kanamycin. Briefly, transformed BL21(DE3) *E. coli* were grown overnight in minimal media (37 °C, 225 RPM) and used to inoculate a main culture flask. Protein expression was induced at an $OD_{600}$ of~0.6 with 1 mM isopropyl β-D-1-thiogalactopyranoside (IPTG) for 5 hours at 37 °C with shaking at 225 RPM. U-[$^{15}$N, $^{13}$C]-labelled 8_4 was expressed in 1× M9 minimal media containing 1 $gL^{-1}$ $^{15}NH_4Cl$, 3 $gL^{-1}$ U-$^{13}$C glucose, 0.1 $gL^{-1}$ $^{15}$N, $^{13}$C ISOGRO, and 12.5 $mgL^{-1}$ Kanamycin. Briefly, transformed BL21(DE3) *E. coli* were grown overnight in minimal media and used to inoculate a main culture flask. Protein expression was induced at an $OD_{600}$ of~0.6 with 1 mM isopropyl β-D-1-thiogalactopyranoside (IPTG) for 5 hours at 37 °C with shaking at 225 RPM.

NMR spectra were processed in NMRPipe and analysed with NMRFAM-SPARKY(*51*, *52*). All NMR experiments were collected on a Bruker Avance III HD 800 MHz spectrometer equipped with a 3 mm TCI cryoprobe at 25°C. Acquisition parameters and pulse sequences used in NMR experiments are listed in table S3. NMR samples were prepared by dialysis into a phosphate-based NMR buffer (20 mM sodium phosphate pH 6.5, 100 mM NaCl, 1 mM EDTA, 0.02% w/v $NaN_3$), followed by addition of 10% $D_2O$. 2D $^{1}$H-$^{15}$N HSQC spectra were collected for 0.66 mM, U-[$^{15}$N, $^{13}$C] labelled apo 8_4 and 0.44 mM U-[$^{15}$N, $^{13}$C]-labelled 8_4 in the presence of 0.62 mM αS: NVGGAVVT peptide (unlabelled, natural abundance). 2D $^{1}$H-$^{15}$N SOFAST HMQC spectra were collected for 60 μM [U-$^{15}$N]-labelled DFASYN013_9 in the absence and presence of 400 μM αS: NVGGAVVT peptide (unlabelled, natural abundance). 2D $^{1}$H-$^{15}$N SOFAST HMQC spectra were collected for 38 μM [U-$^{15}$N]-labelled and DFASYN017_2 in the absence and presence of 240 μM αS: VTAVAQKTVEGAGSIAAATGFV peptide (unlabelled, natural abundance).

Backbone assignment was performed using a 0.44 mM U-[$^{15}$N, $^{13}$C]-labelled 8_4 in the presence of 0.62 mM αS: NVGGAVVT peptide (unlabelled, natural abundance). To assign NH, CO, Cα, and Cβ resonances, standard HSQC-based 3D HNCO, HNCA, HN(CA)CB, CBCA(CO)NH triple resonance experiments were acquired(*53*). Amide to amide NOEs were collected using a 3D SOFAST NOESY-HMQC $H_NH_{Aro}$-$NH_N$ experiment(*54*). For all 3D NMR data acquisition, ~20% non-uniform sampling (NUS) was used with schedules obtained with the hmsIST schedule generator v3.0 with 0.1% tolerance and shuffling enabled (https://scottrobson.org/hmsIST/gensched_new.html)(*55*). RCI-$S^2$ and SSI values were obtained from assigned chemical shifts using TALOS-N(*56*).

Biolayer interferometry

Binding kinetics were measured on an Octet RED96 system (ForteBio) at 25°C with agitation at 1,000 rpm in Octet buffer (10 mM HEPES, 150 mM NaCl, 3 mM EDTA, 0.05% Surfactant P20, 1 mg/mL BSA, pH 7.4). Streptavidin (SA) biosensors were hydrated in Octet buffer for >10 min prior to use. Biotinylated ligands were immobilized onto sensors to a response level of 50-70% saturation, followed by a 60 s baseline and association and dissociation against serial dilutions of analyte.

Enzymatic biotinylation

Avi-tagged constructs were biotinylated *in vitro* using the BirA500 kit (Avidity). IMAC-eluted protein was reacted with BirA ligase according to the manufacturer's instructions (overnight at 4°C or 2–3 h at room temperature). The biotinylated fraction was purified by SEC (Superdex 200 or 75 Increase) to remove free biotin and enzyme.

Alpha synuclein expression, purification and labelling

Wild-type αSyn was expressed and purified as reported previously(*16*). In short, a pT7-7 plasmid transformed into *E.coli* BL21(DE3) Gold cells were cultured in 2× yeast extract tryptone (YT) medium at 37°C and induced with 1 mM IPTG at 28°C. Lysates were clarified and subjected to boiling (80–95°C, 20 min) to precipitate heat-labile proteins. The supernatant was then purified by anion exchange chromatography (HiLoad 26/10 Q Sepharose) followed by SEC on a HiLoad 26/600 Superdex 75 pg column.

Preparation of kinetically trapped oligomers

Kinetically trapped oligomers (KTOs) were prepared according to previous protocols(*37*). Briefly, filtered αSyn (7.5 mg/mL) was incubated at 37°C for 16 h without agitation, removing residual monomer by three rounds of 100 kDa centrifugal filtration, and quantifying by absorbance at 275 nm ($\varepsilon = 5600\ M^{-1}cm^{-1}$).

Microfluidics diffusional sizing

Binding affinities for monomers and oligomers were measured on Fluidity One-M (Fluidic Sciences). Fluorescently labelled binders were incubated with monomers and oligomers at a range of concentrations for 30 min at room temperature. Microfluidic circuits on the Fluidity One-M chip plate were primed with sample buffer immediately before use. Size range setting 2 was selected for all measurements. $K_D$ values were determined by non-linear least squares fitting on the instrument. Fibril binding affinities were measured on smMDS microfluidic devices fabricated using standard soft-lithography techniques and operated as reported previously(*36*, *57*). A negative pressure was applied to the outlet using a Hamilton syringe connected to a syringe pump (Cetoni, neMESYS) to draw the sample and buffer through the hydrophilic-treated channels at a constant flow rate of 100 µL/h. The measurement system was equipped with 488 nm and 647 nm laser lines for excitation and two corresponding single-photon avalanche diode (SPAD) detectors. Diffusion was measured by translocating the confocal volume through the four innermost channels at the mid-height of the device at 20 µm/s. Diffusion profiles were fit to advection diffusion simulations by least squares to recover hydrodynamic radius via the Stokes Einstein relation(*58*), using previously published analysis software(*36*).

Negative-stain electron microscopy

Preformed αSyn fibrils (PFF, 10 µM monomer equivalent) were incubated with gold-conjugated binder (20 µM binder, 10 µL gold conjugate, 50 µL total) for 5 min at room temperature, applied to glow-discharged carbon-coated grids, negatively stained with uranyl acetate, and imaged on a Talos S200X TEM.

Kinetics assay for fibril formation

Lyophilized αSyn was dissolved in 6M GuHCl and subjected to SEC (Superdex 75 Increase) to isolate pure monomers immediately prior to use while exchanging them to the reaction buffer. Aggregation reactions were performed in 1× PBS (pH 7.4) containing 0.01% $NaN_3$ and 50 µM ThT. Seeds were made from aggregating αSyn under non-seeded conditions without the addition of ThT and tip sonicated (1 min, 10% power, 30% cycle) prior to use. Seeds were added as a percentage of the total protein concentration. Samples were loaded onto 96-well plates (Corning 3881) with one 3 mm glass bead (Sigma-Aldrich) per well and their aggregation was monitored by ThT fluorescence (excitation: 440 nm; emission: 480 nm) using FLUOstar plate readers. Reactions were incubated at 37°C with agitation between measurement cycles (200 rpm orbital shaking) and total cycle time matching the minimum cycle time to decrease resting periods between cycles(*59*).

Mammalian cell culture and transfection

Lenti-X 293, HeLa, and HEK293T cells were maintained under standard cell culture conditions in complete DMEM media (Gibco) supplemented with 10 % fetal bovine serum (Gibco), 2 mM L-glutamine (Gibco) and 100 U/ml penicillin/streptomycin (Thermo Fisher). Cells were incubated at 37°C with 5 % $CO_2$ and maintained below ~80 % confluency.

The transient transfection of membrane targeting αSyn NAC domain-mScarlet expressing plasmid followed the Lipofectamine 3000 workflow(*60*). Approximately 4 x $10^4$ cells from each cell line were seeded in a µ-Slide 8 Well plate (Ibidi) 3 days before the experiment. For transfection, 40 µL of Opti-MEM supplied with 1.2 µL of Lipofectamine 3000 was mixed with 80 µL Opti-MEM, 1.6 µg of the plasmid, and 5 µL of P3000 reagent at a 1:1 v/v ratio and incubated at room temperature for 15 min. 7.5 µL of the mixture was added to each well for overnight transfection. For the imaging experiments, doxycycline (500 ng/mL) was added one day after Lipofectamine transfection of the αSyn-mScarlet construct and one day prior to imaging.

Lentivirus transduction and generation of stable binder-EGFP cell line

Lentivirus was produced by transfecting Lenti-X 293 cells with lentiviral transfer vectors, psPAX2 (packaging) and pMD2.G (envelope) at a 3:2:1 ratio using FuGENE HD Transfection Reagent (Promega), Opti-MEM (Thermo Fisher), and the binder-EGFP plasmid. The viral supernatant was collected 48 h after transfection, filtered through a 0.45 μm filter (Millipore) and quantified using a Lenti-X GoStix Plus (Takara Bio).

HeLa cells were seeded into tissue-culture-treated 6-well plates and allowed to reach 80 % confluency. Cells were washed and media changed with 2 mL of lentivirus containing 8 μg/mL polybrene (Tocris Bioscience) before centrifuging at 500 ×*g*, 30 min. Cells were maintained under standard cell culture conditions before selection with 2 μg/mL puromycin (Thermo Fisher).

Confocal imaging

Live cells were stained with Hoechst 33342 (8 μg/mL, 10 min) and imaged on a Leica STELLARIS 5 (HC PL APO CS2 63x, 1.40 oil) with sequential excitation at 405 nm, 488 nm, and 584 nm. Images were acquired at 1024x1024 pixels with the pinhole at 1 Airy unit. Colocalisation between the binder–EGFP and αSyn–mScarlet channels was quantified as the Pearson correlation coefficient using Fiji/ImageJ over > 50 cells.

DAmFRET

HEK293T cells were plated in 96-well plates at 5 x $10^5$ cells per well in 130 μL of medium 18 to 24 h before transfection. Plasmids encoded mEos3.2 tagged αSyn A53T together with a binder or BDFP1.6:1.6 as a negative control. αSyn Type 1 and Type 2 pre-formed fibrils (StressMarq) were diluted twofold to 1 mg/mL and sonicated in a Diagenode Bioruptor Pico water bath sonicator for two cycles of 30 s on and 30 s off. Transfections were assembled in triplicate. The DNA mix consisted of 0.36 μg plasmid, 0.9 μg of sonicated fibrils in treated mixtures, followed by 0.72 μL P3000 reagent, plus Opti-MEM for a final volume of 36 μL. Then, Lipofectamine 3000 was diluted in OptiMEM at a ratio of 0.75 μL Lipofectamine 3000 per 9.25 μL Opti-MEM. This mixture was held at room temperature for 5 min, added to the DNA mix in equal volume, and incubated for a further 20 min. Finally, 20 μL was added per well, giving 0.10 μg of DNA, 0.25 μg fibrils, 0.2 μL P3000, and 0.75 μL Lipofectamine 3000 per well.

After 48 h, cells were detached with 50 μL of TrypLE Express for 5 min at room temperature and collected in a 96-well round bottom plate (Corning). A 50 μL PBS wash was added to the original plate and pooled with the collected cells. Cells were fixed by adding 100 μL of 4% paraformaldehyde for 10 min at room temperature, then pelleted at 500 ×*g* for 5 min. The supernatant was discarded and the pellet was resuspended in 110 μL of PBS containing 10 mM EDTA.

DAmFRET acquisition was conducted in accordance with previous studies(*40*, *41*, *61*, *62*). mEos3.2 was photoconverted using a 405 nm lamp for 3 min 43 sec, and 80 µL of culture per well was analyzed on a Bio-Rad ZE5 cytometer at 1.5 to 1.8 µL/s. Donor fluorescence was excited with a 488 nm laser and collected with a 525/35 detector. FRET was excited with a 488 nm laser and collected with a 593/52 detector. Acceptor fluorescence was excited with a 561 nm laser and collected with a 589/15 detector. BDFP fluorescence was excited with a 640 nm laser and collected with a 670/30 detector. Autofluorescence was measured with a 405 nm laser and collected with a 460/22 detector. Data was analysed in FCS Express. Using forward scatter (FSC) and side scatter (SSC), the gating paradigm removed debris using SSC-A vs FSC-A and selected for single cells using FSC-H vs FSC-A. Expressing cells were gated using Autofluorescence vs Donor fluorescence. Photoconversion efficiency was confirmed using Acceptor fluorescence vs Donor fluorescence. AmFRET was calculated as [FRET-A]/[Acceptor-A] and plotted against Acceptor-A as a proxy for cell expression. A single AmFRET positive gate was applied across all wells.

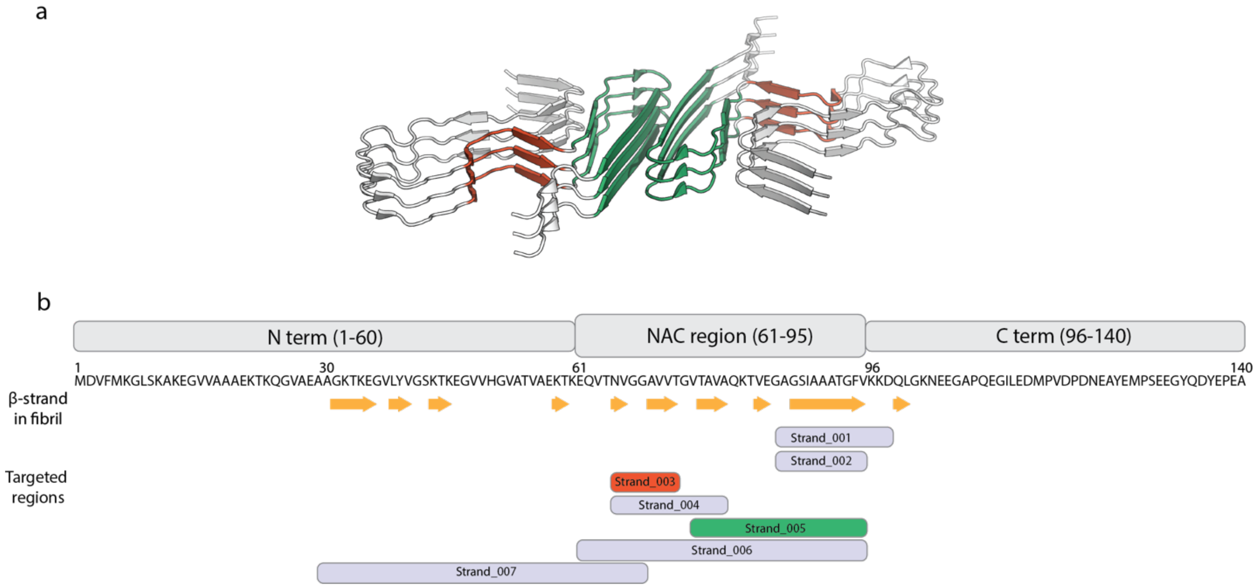


**Fig. S1. Target selection.**

(a) Solid-state NMR structure of αSyn fibrils derived from human Lewy body dementia tissue taken from PDB ID 8FPT. β strand targets which yielded successful binders are coloured red and green. (b) The seven NAC subregions targeted for design, mapped onto the αSyn sequence and the fibril structure.

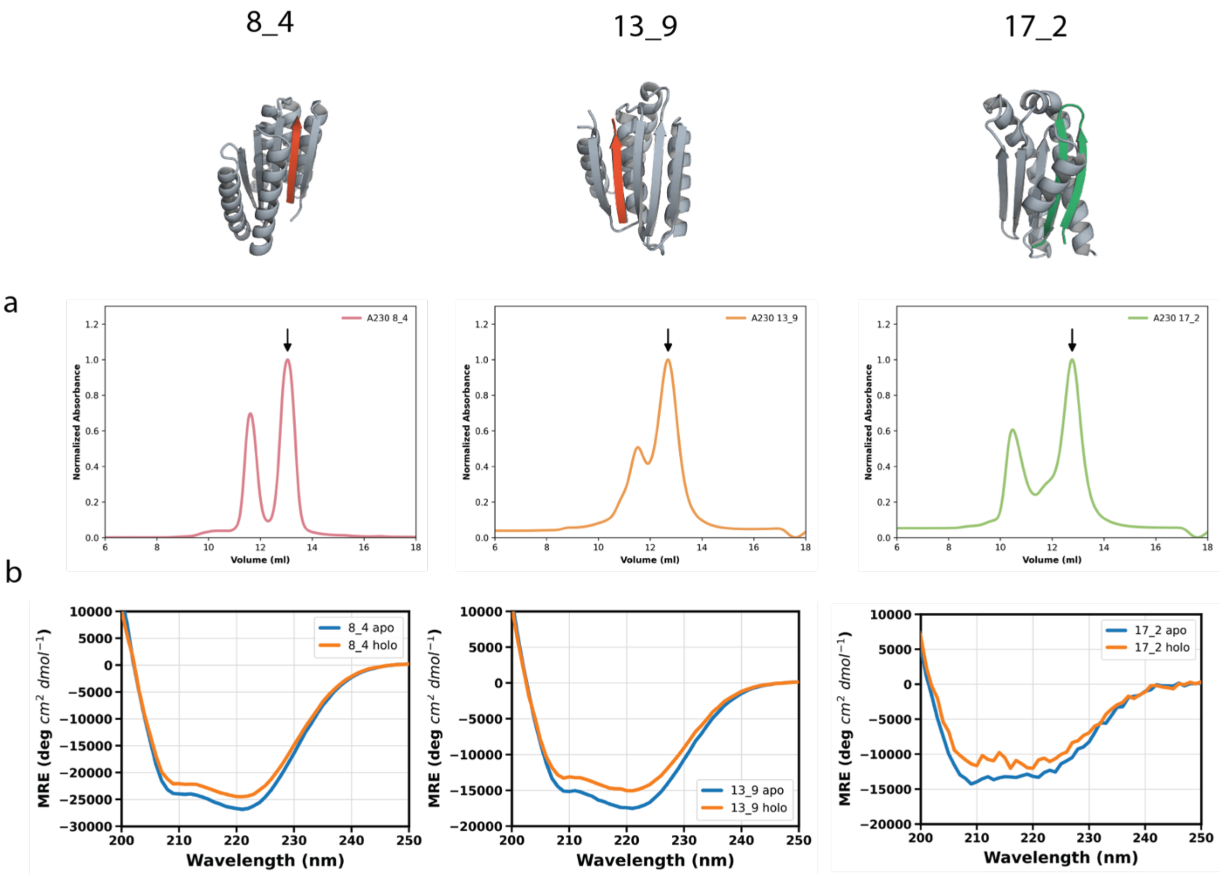


**Fig. S2. Characterisation of the designed binders.**

(a) Size exclusion chromatography traces monitored at 230 nm for the binders. Arrows mark the peak collected and carried forward into all subsequent experiments. (b) Circular dichroism (CD) spectrum of each binder alone (blue) and with their target peptide (orange), measured at 25°C.

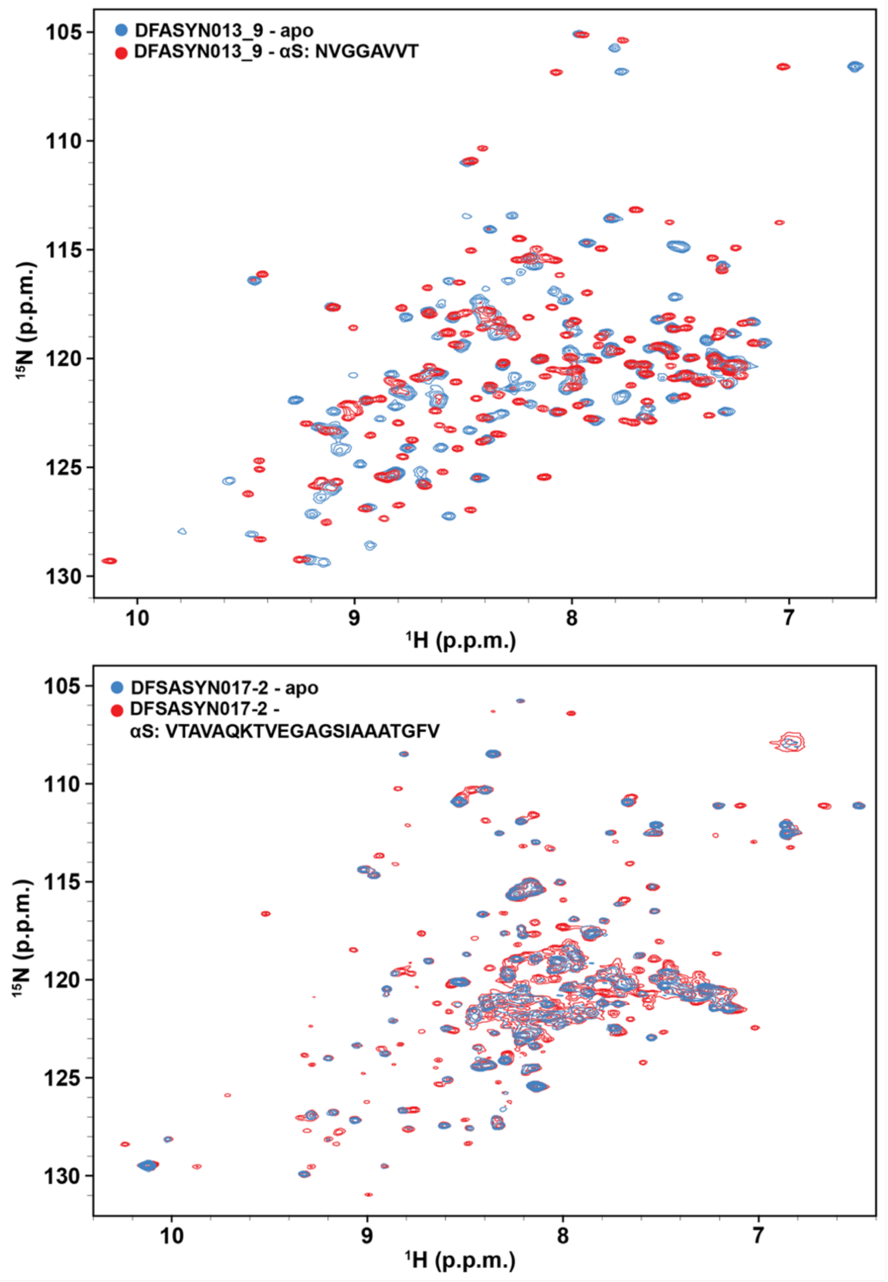


**Fig. S3.**

2D $^{1}$H-$^{15}$N SOFAST HMQC spectra of 60 µM [U-$^{15}$N]-labelled 13_9 (top) and 38 µM [U-$^{15}$N]-labelled 17_2 (bottom) in their free and peptide-bound states (blue and red, respectively) acquired at a $^{1}$H field of 800 MHz at 25 °C. The bound states (red) contain molar excess of unlabelled, natural abundance peptide: 400 µM αS: NVGGAVVT (with 13_9) or 240 µM αSyn: VTAVAQKTVEGAGSIAAATGFV (with 17_2).

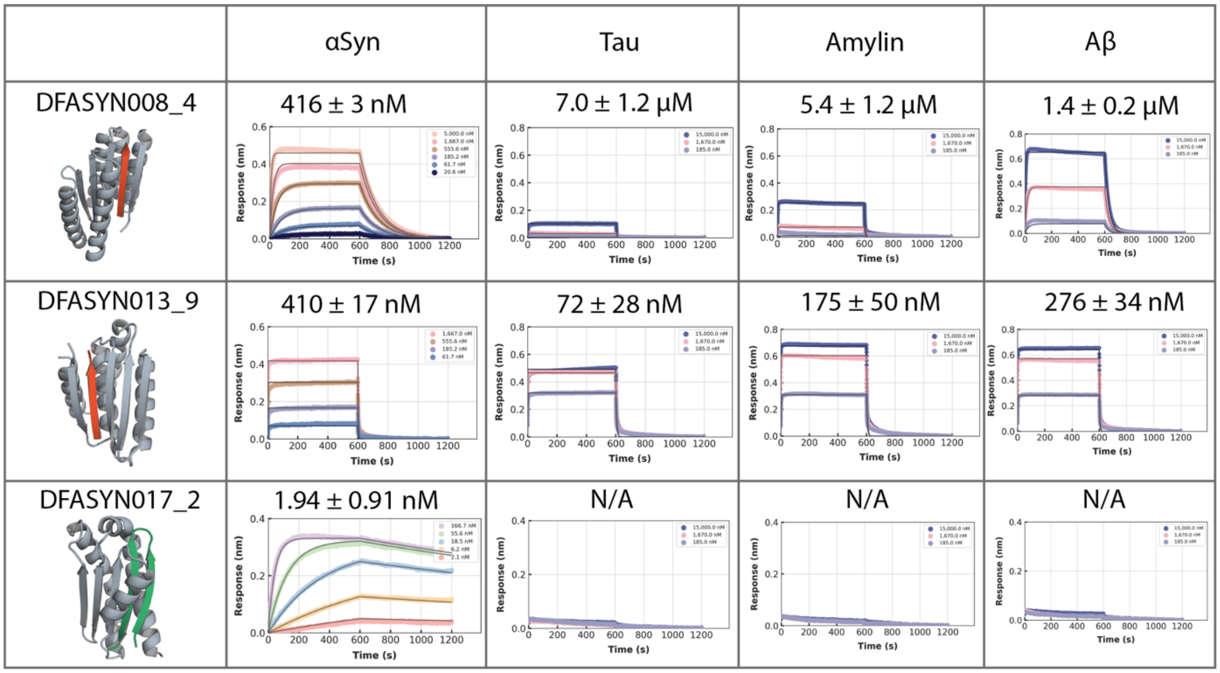


**Fig. S4. Selectivity of the matured binders against related amyloidogenic cores.**

BLI of 8_4, 13_9 and 17_2 against ubiquitin fused peptides representing the aggregation cores of αSyn, tau, amylin and amyloid β (Supplementary Table 2), with the designed structure of each binder shown at left and the fitted dissociation constant above each sensorgram.

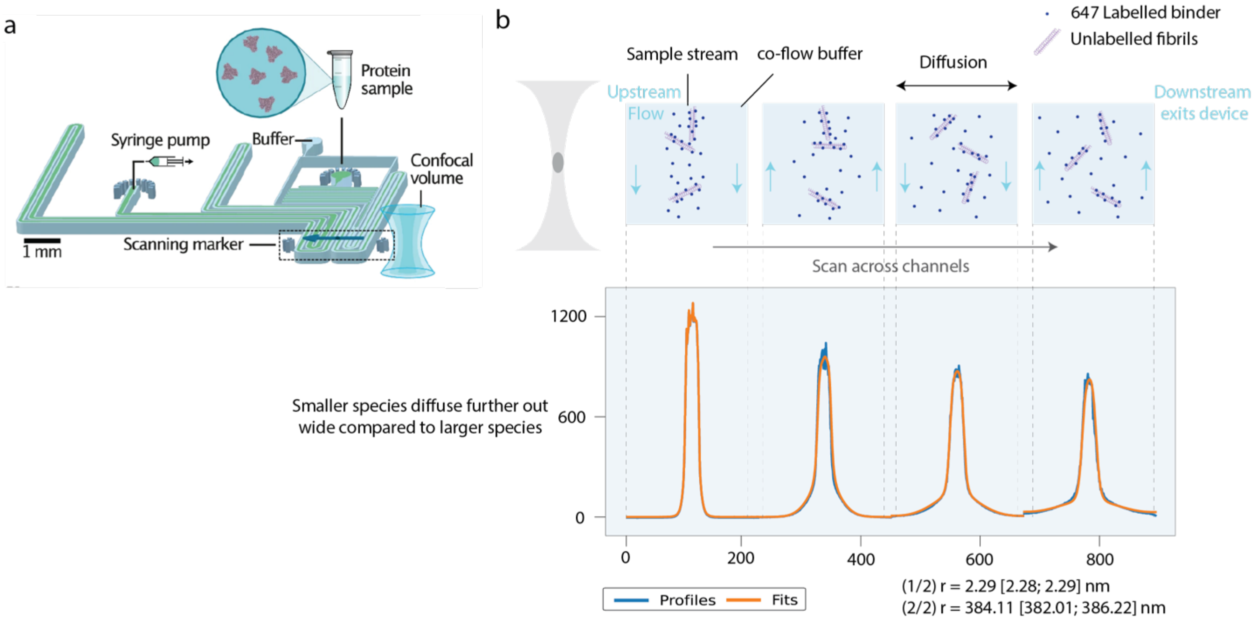


**Fig. S5. Principle of single-molecule microfluidic diffusional sizing (smMDS).**

(a) smMDS microfluidic chip design. Reproduced with permission from *Krainer et al*.(*36*) (b) Schematic of the operating principle of smMDS. Species are diffused laterally and the fluorescence signal can be fitted.

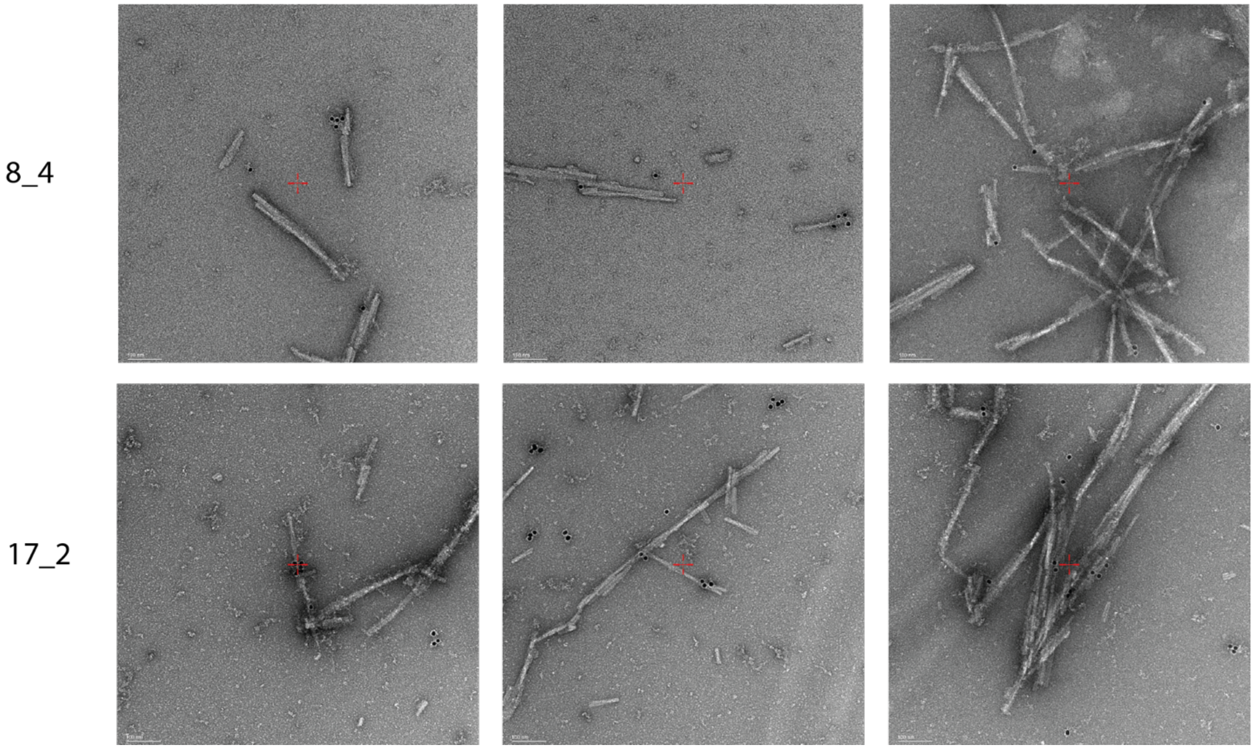


**Fig. S6. Negative stain TEM images showing gold-conjugated binders binding to fibrils.**

8_4 shows preferential binding towards the fibril ends whereas 17_2 showed preferential binding to the fibril surfaces and low molecular weight aggregates.

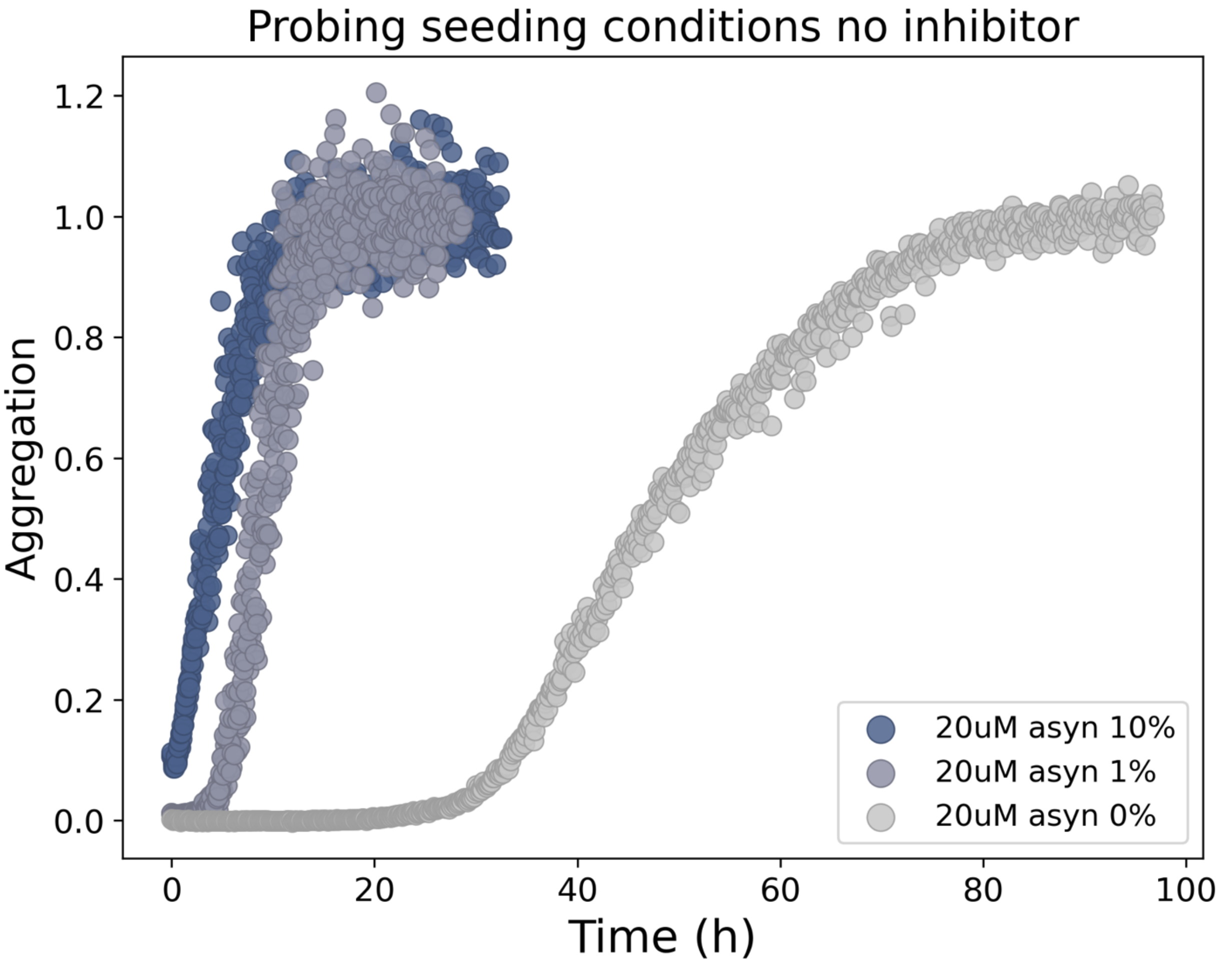


**Fig. S7. Probing sufficient seeding conditions for αSyn inhibition without inhibitor.**

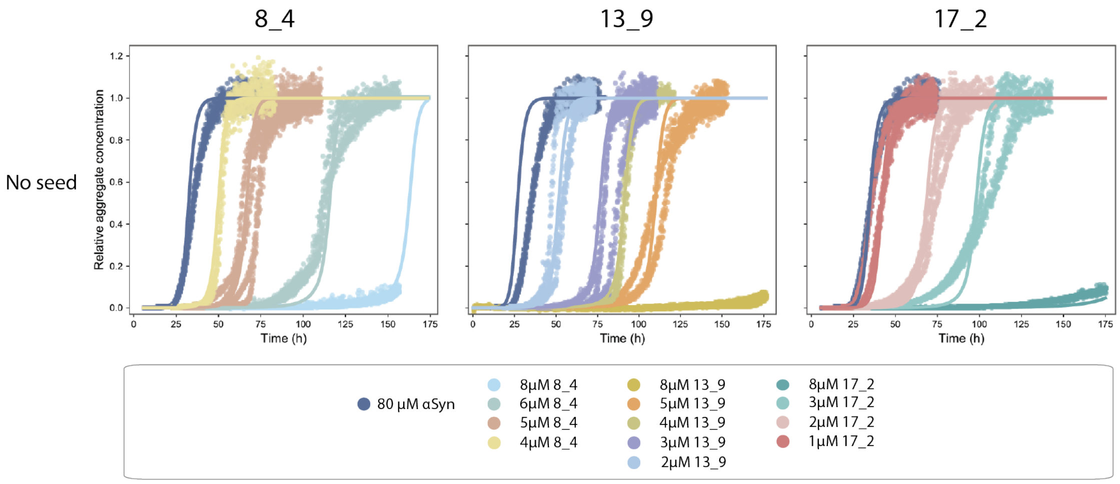


**Fig. S8. Titrating binders at no seed conditions shows no significant impact on primary nucleation, fitted using the same model as high seed and low seed conditions.**

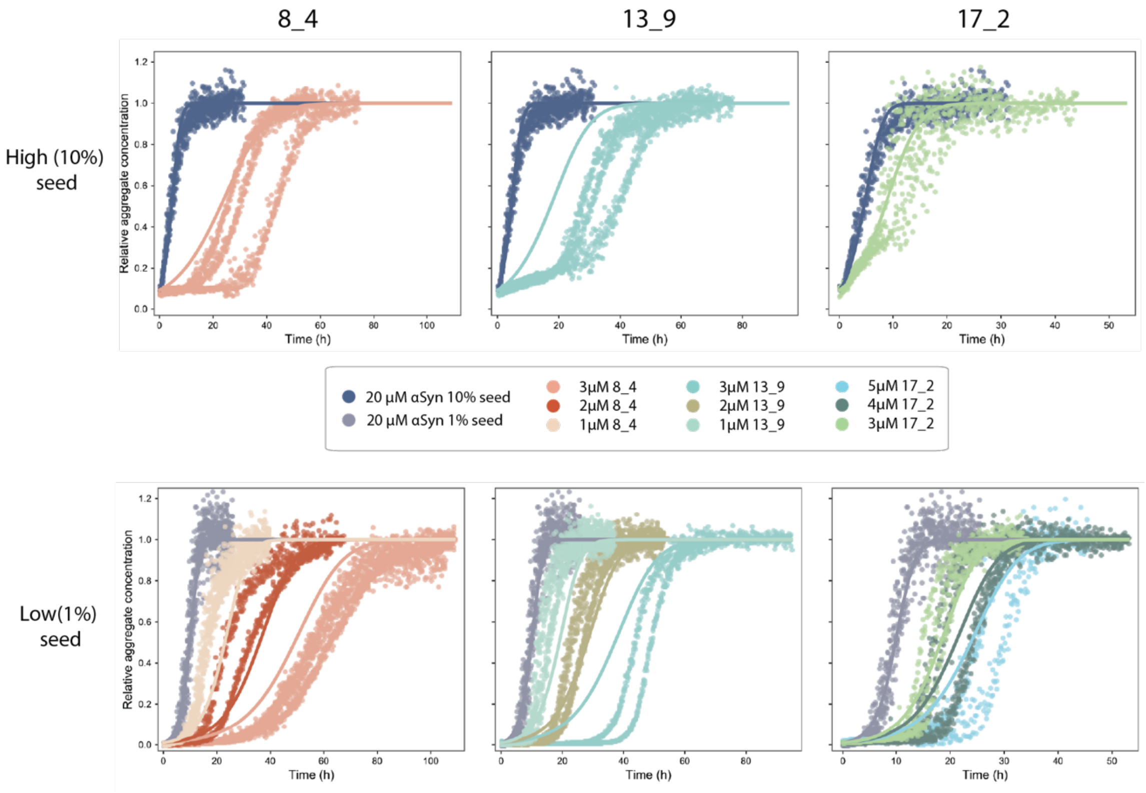


**Fig. S9. Bad fits excluding the depletion model, showing that binders preferentially bind aggregates.**

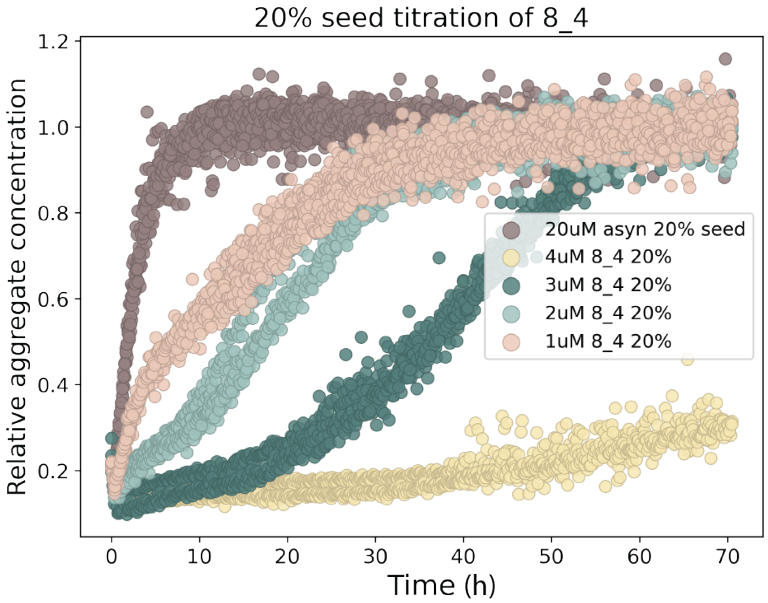


**Fig. S10. Titration of 8_4 at high (20%) seed conditions.**

Substoichiometric ratios of 8_4 inhibited the aggregation strongly, showing strong inhibition ability through interaction with fibril ends.

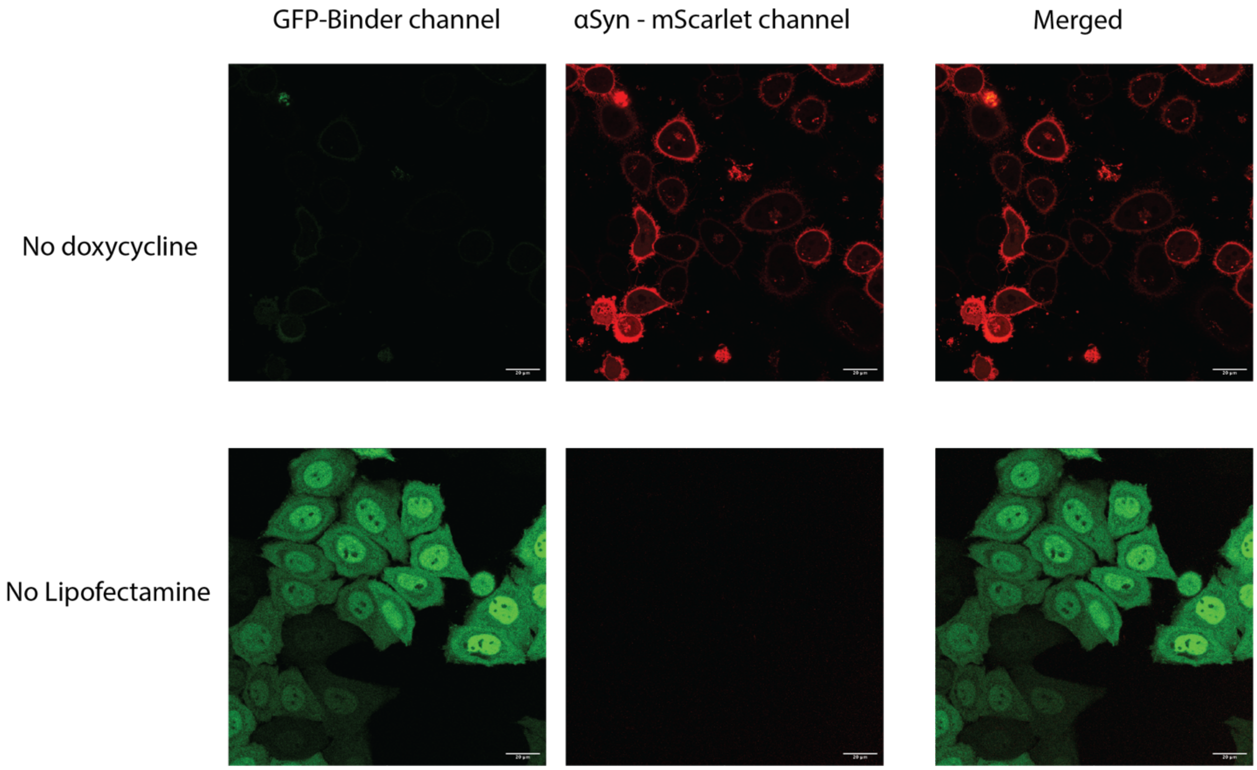


**Fig. S11. Cell controls.**

Confocal images for the no doxycycline and no lipofectamine conditions, confirming the dependence of signal on both partners.

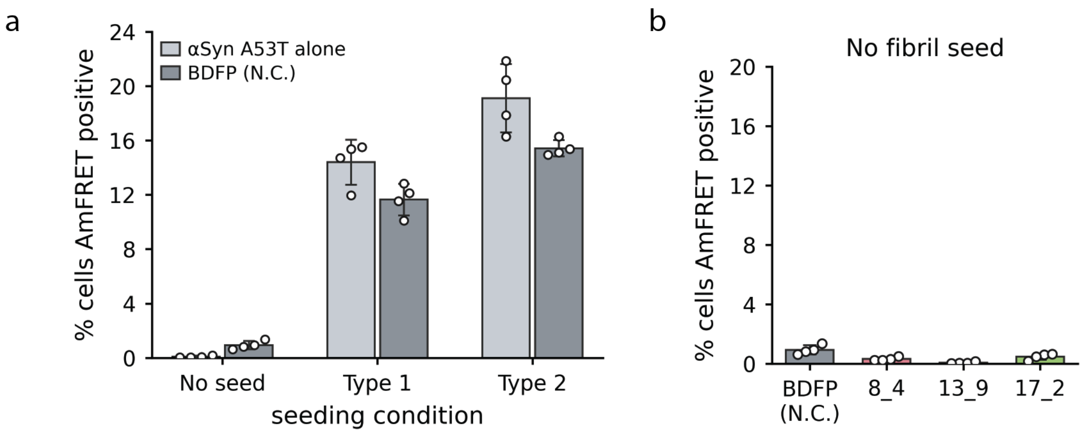


**Fig. S12.**

(a) αSyn A53T alone versus BDFP for each seeding condition. (b) All constructs without fibril seed, on the Figure 5d axis.

| Metric | Initial designs | After partial diffusion | Change (%) |
|---|---|---|---|
| pLDDT | 94.92 | 95.57 | +0.65 |
| Mean pAE (Å) | 3.4 | 2.9 | -15.11 |
| Max pAE (Å) | 19.8 | 13.1 | -33.69 |
| Cα-RMSD (Å) | 0.71 | 0.62 | -12.40 |

**Table S1. Improvements in ColabFold metrics after partial diffusion**

| Peptide target | Amino acid sequence |
|---|---|
| αSyn | TKEQVTNVGGAVVTGVTAVAQKTVEGAGSIAAATGFVKKD |
| Tau | GGSVQIVYKP |
| Amylin | NFGAILSST |
| Aβ | SNKGAIIGLMV |

**Table S2. Peptide ligands for BLI experiments**

| Experiment (Bruker pulse sequence) | Nucleus | Sweep width (Hz) | Number of points (TD) | Acquisition time (ms) | Relaxation delay (s) | Mixing time (s) |
|---|---|---|---|---|---|---|
| 3D HNCO (hncogp3d) | $^{1}H$ | 12820.5 | 2048 | 79.9 | 1 | N/A |
| | $^{15}N$ | 2513.6 | 48 | 9.5 | | |
| | $^{13}CO$ | 2817.1 | 128 | 22.7 | | |
| 3D HNCA (hncagp3d) | $^{1}H$ | 12820.5 | 2048 | 79.9 | 1 | N/A |
| | $^{15}N$ | 2513.6 | 48 | 9.5 | | |
| | $^{13}C_{\alpha}$ | 6036 | 128 | 10.6 | | |
| 3D HNCACB (hncacbgp3d) | $^{1}H$ | 12820.5 | 2048 | 79.9 | 1 | N/A |
| | $^{15}N$ | 2513.6 | 48 | 9.5 | | |
| | $^{13}C_{\alpha,\beta}$ | 16095.8 | 160 | 5 | | |
| 3D CBCACONH (cbcaconhgp3d) | $^{1}H$ | 12820.5 | 2048 | 79.9 | 1 | N/A |
| | $^{15}N$ | 2513.6 | 48 | 9.5 | | |
| | $^{13}C_{\alpha,\beta}$ | 16095.8 | 160 | 5 | | |
| 3D SOFAST NOESY-HMQC $H_NH_{Aro}$-$NH_N$ (sfnoesyhmqc3d.HnHa-NHn) | $^{1}H$ | 12820.5 | 2048 | 79.9 | 0.2 | 0.3 |
| | $^{15}N$ | 2513.6 | 64 | 12.7 | | |
| | $^{1}H$ | 4400.7 | 144 | 16.4 | | |
| 2D $^{1}H$-$^{15}N$ SOFAST HMQC (IBS_SOFAST.x) | $^{1}H$ | 12820.5 | 2048 | 79.9 | 0.2 | N/A |
| | $^{15}N$ | 2756.9 | 256 | 46.4 | | |
| 2D $^{1}H$-$^{15}N$ HSQC (hsqcetf3gpsi) | $^{1}H$ | 12820.5 | 2048 | 79.9 | 1 | N/A |
| | $^{15}N$ | 2513.6 | 512 | 101.8 | | |
| | $^{15}N$ | 2756.87 | 512 | 92.9 | | |

**Table S3. NMR acquisition parameters and pulse sequences.**

| Binder | $K_{IE}$ (M) | $K_{IS}$ (M) | $K_{IE, on}$ ($M^{-1}$ $h^{-1}$) | $K_{IS, on}$ ($M^{-1}$ $h^{-1}$) | $SE_{Low}$ | $SE_{High}$ |
|---|---|---|---|---|---|---|
| **8_4** | $9.76\times10^{-10}$ | $3.51\times10^{-6}$ | $9.11\times10^{5}$ | $1.02\times10^{8}$ | 1.15 | 0.350 |
| **13_9** | $1.43\times10^{-8}$ | $3.61\times10^{-6}$ | $7.26\times10^{5}$ | $2.13\times10^{6}$ | 1.15 | 0.350 |
| **17_2** | $3.17\times10^{-5}$ | $1.27\times10^{-7}$ | $3.40\times10^{8}$ | $2.67\times10^{6}$ | 1.15 | 0.680 |

**Table S4. Global kinetic fitting parameters for the three binders.**

For all fittings, $k_2$ = 0.338 $h^{-1}M^{-0.228}$, $n_2$ = 0.228, and $k_+$ = 1.73 × $10^5$ $h^{-1}M^{-1}$, and the average seed length $L_0$ = 15.5 monomer units. Values are best fit ± 95% confidence interval from 3 independent replicates.

| | |
|---|---|
| $k_2$ | Secondary nucleation rate constant ($M^{-0.228}$ $h^{-1}$) |
| $n_2$ | Secondary nucleation reaction order |
| $k_+$ | Elongation rate constant ($M^{-1}$ $h^{-1}$) |
| $K_{IE}$ | Dissociation constant of the binder for growth competent fibril ends (M) |
| $K_{IS}$ | Dissociation constant of the binder for catalytic surface sites (M) |
| $K_{IE, on}$ | Association rate constant of the binder for fibril ends ($M^{-1}$ $h^{-1}$) |
| $K_{IS, on}$ | Association rate constant of the binder for catalytic surface sites ($M^{-1}$ $h^{-1}$) |
| $SE_{Low}$ | Scaling factor on the added seed mass at 1% seed |
| $SE_{High}$ | Scaling factor on the added seed mass at 10% seed |
